# Classical and controlled auditory mismatch responses to multiple physical deviances in anaesthetised and conscious mice

**DOI:** 10.1101/831016

**Authors:** Jamie A. O’Reilly, Bernard A. Conway

## Abstract

Human mismatch negativity (MMN) is modelled in rodents and other non-human species to examine its underlying neurological mechanisms, primarily described in terms of deviance-detection and adaptation. Using the mouse model, we aim to elucidate subtle dependencies between the mismatch response (MMR) and different physical properties of sound. Epidural field potentials were recorded from urethane-anaesthetised and conscious mice during oddball and many-standards control paradigms; with stimuli varying in duration, frequency, intensity, and inter-stimulus interval. Resulting auditory evoked potentials, classical MMR (*oddball – standard*), and controlled MMR (*oddball – control*) waveforms were analysed. Stimulus duration correlated with stimulus-off response peak latency (p < 0.0001). Frequency (p < 0.0001), intensity (p < 0.0001), and inter-stimulus interval (p < 0.0001) correlated with stimulu-son *N*1 and *P*1 (conscious only) peak amplitudes. These relationships were instrumental in shaping classical MMR morphology in both anaesthetised and conscious animals, suggesting these waveforms reflect modification of normal auditory processing by different physical properties of stimuli. Controlled MMR waveforms appeared to exhibit habituation to auditory stimulation over time, which was equally observed in response to oddball and standard stimuli. These observations are not consistent with the mechanisms thought to underlie human MMN, which currently do not address differences due to specific physical features of auditory deviance. Thus, no evidence was found to objectively support the deviance-detection or adaptation hypotheses of MMN in relation to anaesthetised or conscious mice.

## 1. Introduction

Mismatch negativity (MMN) is widely believed to reflect pre-attentive sensory-memory. It is observed in response to an oddball paradigm, in which many identical standard stimuli are infrequently replaced with deviant oddball stimuli. This study concerns the auditory modality, as does the majority of literature on MMN. Oddball stimuli may deviate from the standard in any discriminable feature of sound (Pakarinen et al., 2007). The auditory-evoked potential (AEP) from the standard is conventionally subtracted from that of the oddball, generating a difference waveform that constitutes the MMN. This may be referred to as the classical approach. Interestingly, MMN is observed from patients in anaesthetised and comatose states (Fischer et al., 1999, 2000; Kane et al., 1996), suggesting that the mechanisms responsible for its generation are not reliant on consciousness. The prevailing theory is that a mismatch between novel auditory input and a memory-representation, or predictive model, of recently experienced auditory inputs, triggers updating of this model, thereby subconsciously capturing the attention. This manifests in scalp-recorded EEG as the MMN component (Winkler et al., 1996). This interpretation has remained largely unchanged since its rationale was first proposed (Näätänen et al., 1978), and aligns comfortably within the theory of predictive coding (Wacongne et al., 2012). Some researchers have cited a lack of cellular neurophysiological evidence for this interpretation, however, instead referring to known properties of sensory neurons, such as stimulus-specific adaptation (SSA), as possible mechanisms of MMN generation (Jääskeläinen et al., 2004; May and Tiitinen, 2010; Ulanovsky et al., 2003). These two propositions, which are referred to as the memory and adaptation hypotheses of MMN, are not necessarily mutually exclusive, as some researchers have pointed out (Garrido et al., 2009); although the extent of their relative contributions remains unknown, and a complete description of the underlying mechanism(s) of MMN generation remains elusive.

Our foremost concern with the proposed mechanisms is that neither accounts for differences in MMN that emerge due to changes in specific physical features of sound. Mismatch negativity waveforms in response to oddball stimuli varying in duration, frequency, intensity, and other parameters are thought to display unique signatures; potentially offering clinical utility in measuring discrete neurophysiological functions (Näätänen et al., 2004; Pakarinen et al., 2007). Source localisation studies have indicated that generators of MMN to duration, frequency, and intensity oddball stimuli reside in separate compartments of the auditory cortex (Frodl-Bauch et al., 1997; Molholm et al., 2004; Rosburg, 2003). Moreover, there is some evidence suggesting that specific deficits may reflect different aspects of neuropsychiatric disease. For instance, it is generally accepted that duration MMN deficits are more prominent in the early stages of schizophrenia, whereas frequency deficits, if present, are more likely to appear following prodromal phase (Atkinson et al., 2012; Erickson et al., 2016; Michie et al., 2000; Todd et al., 2008; Umbricht and Krljes, 2005). Taken together, this evidence suggests that at least partially distinct networks are responsible for generating duration and frequency MMN. This concept may logically be extended to other physical dimensions of auditory deviance. It may also be noted that SSA is almost exclusively studied in response to stimulus frequency changes, and there is little evidence, to our knowledge, that demonstrates duration- or intensity-specific adaptation; some findings have actually contravened intensity-induced SSA (Duque et al., 2016). Thus, the two most prominent hypotheses of MMN may be over-generalized to completely describe the observed neurophysiological phenomena.

The mouse is a desirable model species for more invasive examination of this clinically relevant biomarker. There are several issues to address concerning pre-clinical translation. Differences in neuroanatomy and electrophysiology recording techniques can produce substantially different AEP waveforms in terms of latency, polarity, and amplitude of different characteristic peaks. Hence when referring to animal studies the term mismatch response (MMR) may be adopted, opposed to MMN, to reflect a reduced emphasis on latency and polarity (Harms et al., 2016). The literature is somewhat conflicted regarding the mouse MMR. An initial study highlighted duration MMR in mice as potentially analogous to human duration MMN, although concluded that frequency MMR was not representative of the human frequency MMN (Umbricht et al., 2005). Nevertheless, researchers have generally continued to favour investigation of frequency oddball paradigms in mice, preserving the established interpretation from human MMN (Ehrlichman et al., 2009, 2008; Featherstone et al., 2015, 2013). Some studies indicate that strain plays an important role in AEP morphology (Ehlers and Somes, 2002; Siegel et al., 2003). This may be important considering that different mouse strains have been used in MMR studies, which could possibly account for variable results. Furthermore, differences in electrophysiology recording protocol, oddball paradigms, and interpretation of control waveforms also obscure direct comparison of findings. Overall, current evidence for a mouse MMR analogous to the human MMN may be described as promising but inconclusive (Featherstone et al., 2018).

Several different control sequences have been employed in rodent MMR studies. One of the most widely accepted is the *many-standards* control paradigm, in which multiple physically-distinct stimuli, including those used in the oddball paradigm, are presented in pseudo-random order at the same rate as deviants in the oddball paradigm (Harms et al., 2014; Jacobsen and Schröger, 2003; Wiens et al., 2019). This is designed to control for oddball presentation rate within an irregular sensory-memory trace. Comparison between AEP waveforms produced by the same stimulus in oddball and many-standards paradigms may therefore be used to discriminate between memory- and adaptation-based MMR. In the *flip-flop* control, assignment of standard and oddball stimuli is alternated in two successive presentations of the oddball paradigm (Harms et al., 2014). The *balanced* oddball paradigm includes two oddball stimuli deviating in the same physical dimension (one increasing and one decreasing) interspersed between standards. This enables a comparison of two separate MMR waveforms produced by subtracting the standard AEP from the two different oddball AEPs. In contrast, the flip-flop control would require four paradigms to examine two difference waveforms relative to the same standard. Given the scope of physical parameters examined, the balanced oddball paradigm and many-standards control are used in the present study.

Considering the inclusion of control paradigms, a distinction may be made between classical (*oddball – standard*) and more stringent controlled (*oddball – control*) methods of MMR computation in the preclinical literature. This aims to dissociate between memory and adaptation contributions to MMR generation, and control for different physical levels of stimuli. However, the majority of clinical literature still employs the classical method (Erickson et al., 2016). This raises questions over the translational reliability of rodent MMR for representing human MMN, motivating a closer inspection of both conventional and more rigorously controlled methods of extracting MMR waveforms.

We hypothesise that different physical deviances may influence MMR wave-forms recorded from the mouse auditory cortex in a feature-specific manner. This study aims to test this hypothesis in urethane-anaesthetised and conscious mice using duration, frequency, intensity, and inter-stimulus-interval (ISI) oddball paradigms. Incorporating many-standards control paradigms, *memory*- versus *adaptation*-based generators are assessed, while characterising the effects of these physical parameters on the resulting mouse AEP and MMR waveforms.

## 2. Calculation

Oddball and many-standards paradigms using each feature of sound were presented sequentially, as described in the Materials and Methods. Habituation of the auditory system in response to continued stimulation, characterised by a progressive decline in AEP amplitudes, is a well-established phenomenon in auditory neurophysiology (Picton et al., 1976). Evoked potentials elicited in consecutive paradigms appeared to demonstrate this phenomenon. Due to this, four waveform comparisons were evaluated, as outlined in Figure 1. *Classical MMR* was generated by subtracting the standard AEP (STD_*OD*_) from the oddball AEP (DEV_*OD*_). These stimuli were physically distinct but presented during the same time interval. Therefore this waveform may reflect sensory-memory disruption, differential adaptation, and/or physical sensitivity of the auditory system. *Controlled MMR* was evaluated by subtracting physically identical control AEPs (DEV_*CTR*_) from their respective oddball AEP; whereas *Controlled Standard* waveforms were computed by subtracting the AEP from standard stimuli presented in oddball and many-standards control paradigms (STD_*CTR*_). In both cases, stimuli were physically identical but presented in two separate time intervals, because paradigms were sequentially ordered. The controlled MMR waveform could reflect oddball-induced sensory memory violation and/or habituation of the auditory response over time; whereas the controlled standard waveform could contain differential adaptation and auditory habituation. This controlled standard differs from previous studies that subtracted the standard AEP from the control AEP, based on the presumption of subtracting a more adapted response from a less adapted response (Harms et al., 2014; Parras et al., 2017). Auditory habituation appeared to be a far more plausible explanation for the changes in AEP amplitudes observed between oddball and many-standards control paradigms in the present study (Figure 4 and Figure 7); therefore we opted to subtract more habituated waveforms from less habituated waveforms. Reversing the order of subtraction would simply invert the polarity of resulting waveforms. To account for physical differences in the oddball paradigm and habituation differences between oddball and control paradigms, controlled MMR and controlled standard waveforms were compared. Termed here as *Normalized MMR*, this analysis should reveal any oddball-induced memory response dissociated from physical sensitivities and auditory habituation; theoretically this could also reflect differential adaptation, although our analysis does not support this.

**Figure 1:**
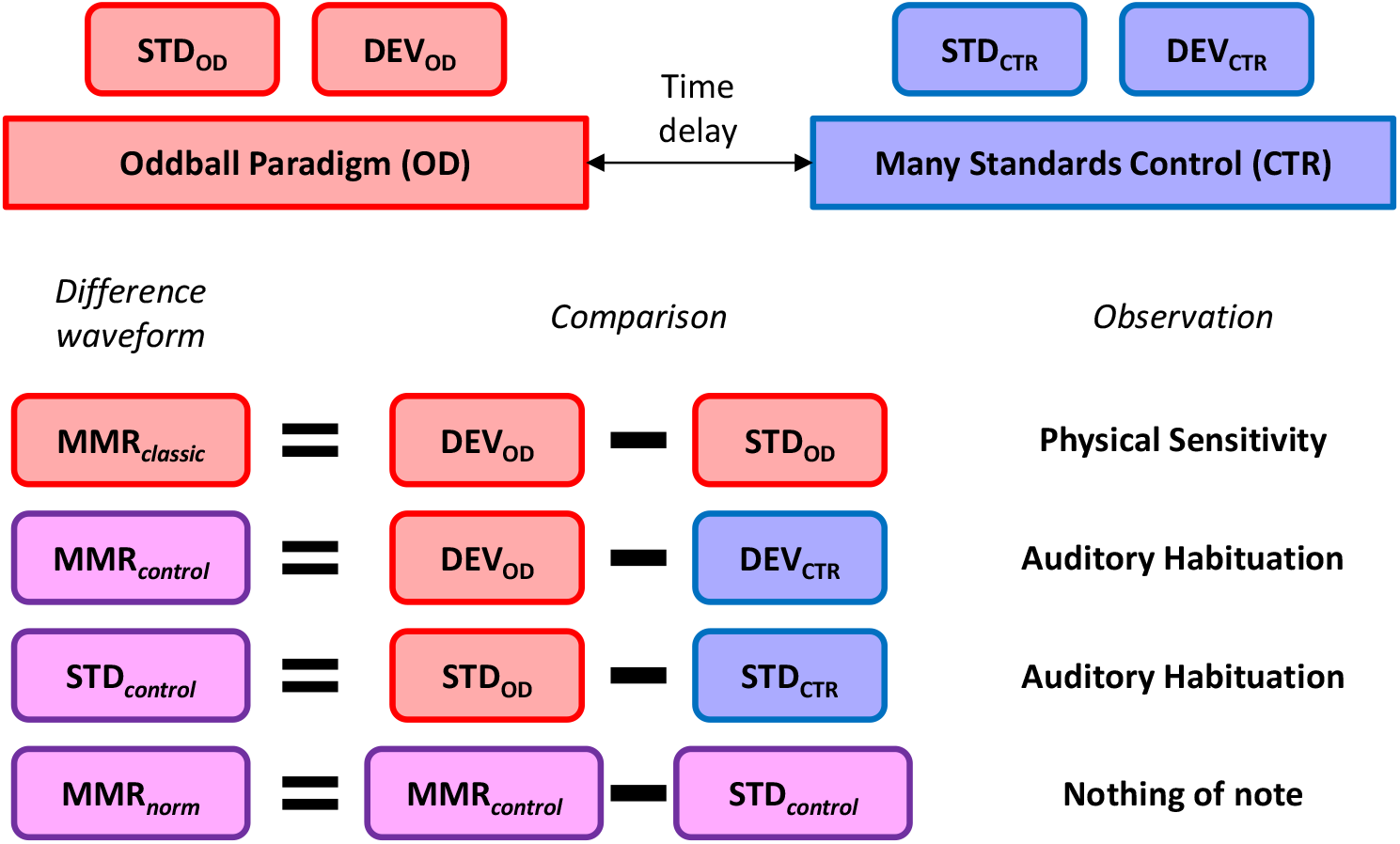
Comparisons between waveforms elicited by oddball and many-standards control paradigms. These are designed to identify any differences between standard, oddball, and control conditions. Observations from these analyses summarily indicated physical sensitivity and habituation of the auditory response.

## 3. Results

### 3.1 Anaesthetised Group

The AEP waveforms from duration, frequency, and intensity many-standards paradigms in urethane-anaesthetised mice are plotted in Figure 2. The effects of these physical parameters are pronounced. Stimulus duration was directly responsible for *P*_*off*_ peak latency [F_9,130_ = 9.50, p < 0.0001]. Stimulus frequency had a significant relationship with *N*1 peak amplitude [F_9,130_ = 5.73, p < 0.0001]. However, stimulus frequency did not significantly affect *P*_*off*_ peak amplitude [F_9,130_ = 1.45, p = 0.175]. On the other hand, stimulus intensity significantly influenced *N*1 [F_9,130_ = 4.91, p < 0.0001] and *P*_*off*_ [F_9,130_ = 5.01, p < 0.0001] peak amplitudes. Interestingly, lower frequencies (<3.75 kHz) appeared to cause a small positive peak amplitude deflection immediately following the *N*1 peak. These stimuli are likely nearing the inaudible frequency range for these animals (Ison et al., 2007). In contrast, low intensity 10 kHz stimuli produced a small amplitude *N*1 without being followed by a positive amplitude deflection. These observations led to an adjustment of stimulus ranges in the experiment with conscious mice, as described in the Materials and Methods.

**Figure 2:**
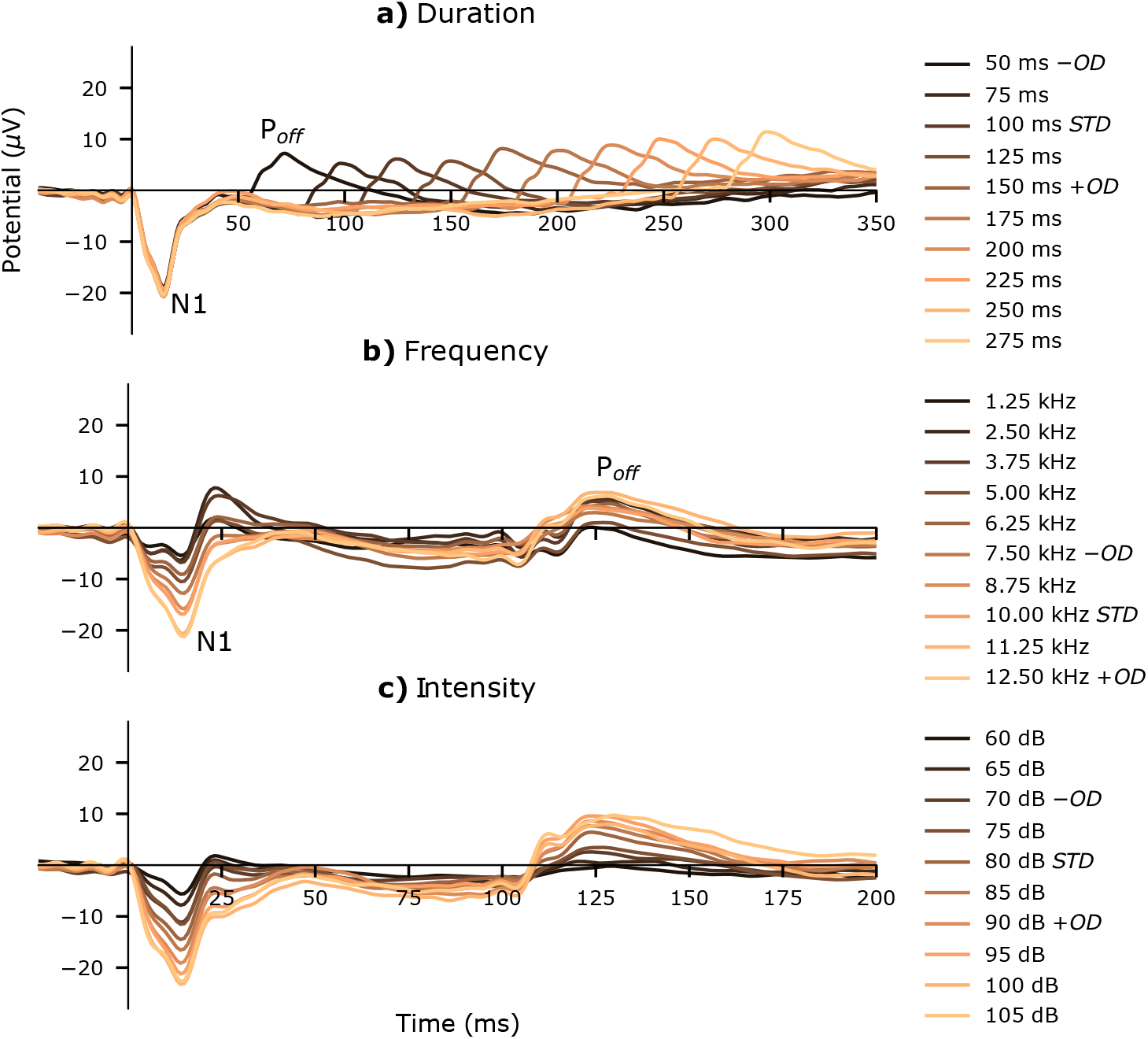
AEP waveforms from many-standards paradigms in urethane-anaesthetised mice. This is grand-average data (n = 14). a) Duration-varying stimuli. b) Frequency-varying stimuli. c) Intensity-varying stimuli. Two components of the AEP are identified; a negative amplitude stimulus onset response (*N*1) and positive amplitude stimulus offset response (*P*_*off*_). Corresponding increasing oddball (+OD), decreasing oddball (−OD), and standard (STD) stimuli from respective oddball paradigms are identified. Note the timescale for duration-varying many-standards control differs from frequency and intensity paradigms.

The classical MMR difference waveforms from duration, frequency and intensity oddball paradigms in urethane-anaesthetised mice are plotted in Figure 3. It is evident from each of these waveforms that the respective physical properties of sound are instrumental in determining MMR morphology. Duration MMR waveforms are caused by *P*_*off*_ potentials occurring at different latencies; positive and negative peak amplitudes corresponding to oddball and standard stimuli offset responses, respectively. These produced regions of significant difference, as determined by statistical analysis, which are annotated in the right-hand panel of Figure 3a. Frequency MMR waveforms are predominantly influenced by *N*1 amplitude modulation by stimulus frequency; for this reason, increasing- and decreasing-frequency oddball stimuli produce opposite polarity deflections over the *N*1 latency range. Similarly, intensity MMR waveforms are generated by *N*1 and *P*_*off*_ amplitude modulation, and increasing- and decreasing-intensity oddballs are observed to produce opposite polarity deflections across *N*1 and *P*_*off*_ latency ranges. Frequency and intensity oddball paradigms generated some regions of statistically significant difference within this measurement window which can be attributed to the observed physical sensitivities; previously observed long-latency potentials from frequency and increasing intensity oddball stimuli are not displayed here (O’Reilly, 2019a). Controlled MMR waveforms are plotted in Figure 4 There are marginal differences between oddball and many-standards paradigm AEP waveforms elicited by physically identical stimuli; reflected in both oddball- and standard-minus-control difference waveforms. These minor differences were mainly concentrated at the *N*1 peak latency, indicative of onset response habituation to repeated stimulation.

**Figure 3:**
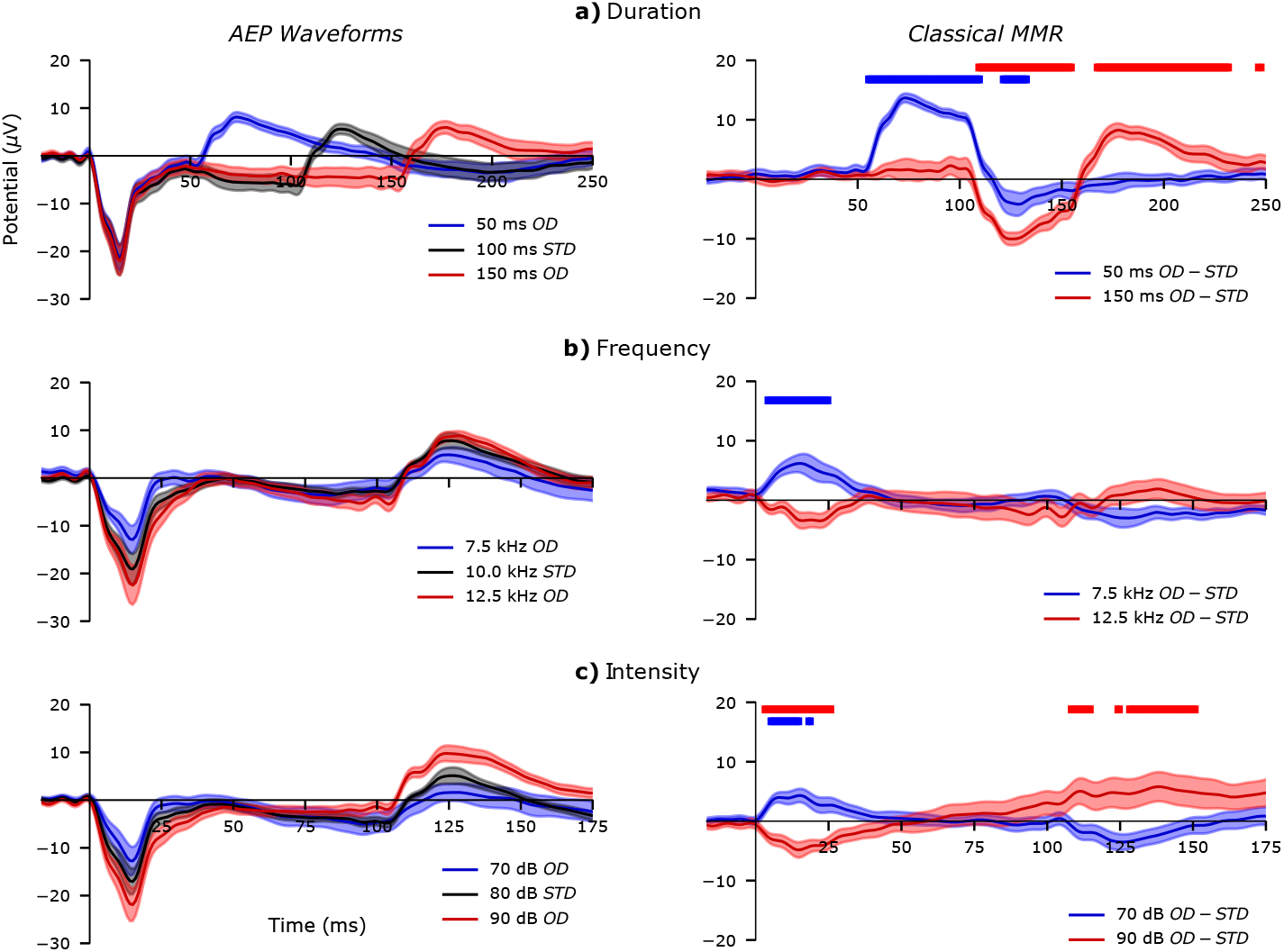
AEP (left) and classical MMR (right) waveforms from oddball paradigms in urethane-anaesthetised mice. This is grand-average data (n = 14). a) Duration oddballs minus 100 ms standard (50 ms OD – STD and 150 ms OD – STD). b) Frequency oddballs minus 10 kHz standard (7.5 kHz OD – STD and 12.5 kHz OD – STD). c) Intensity oddballs minus 80 dB standard (70 dB OD – STD and 90 dB OD – STD). Latencies with statistically significance difference between OD and STD waveforms (p < 0.05) are denoted in the MMR plots with solid bars, coloured to match the corresponding OD waveform. Note the different timescale for duration oddball waveforms. Shaded regions represent sem.

**Figure 4:**
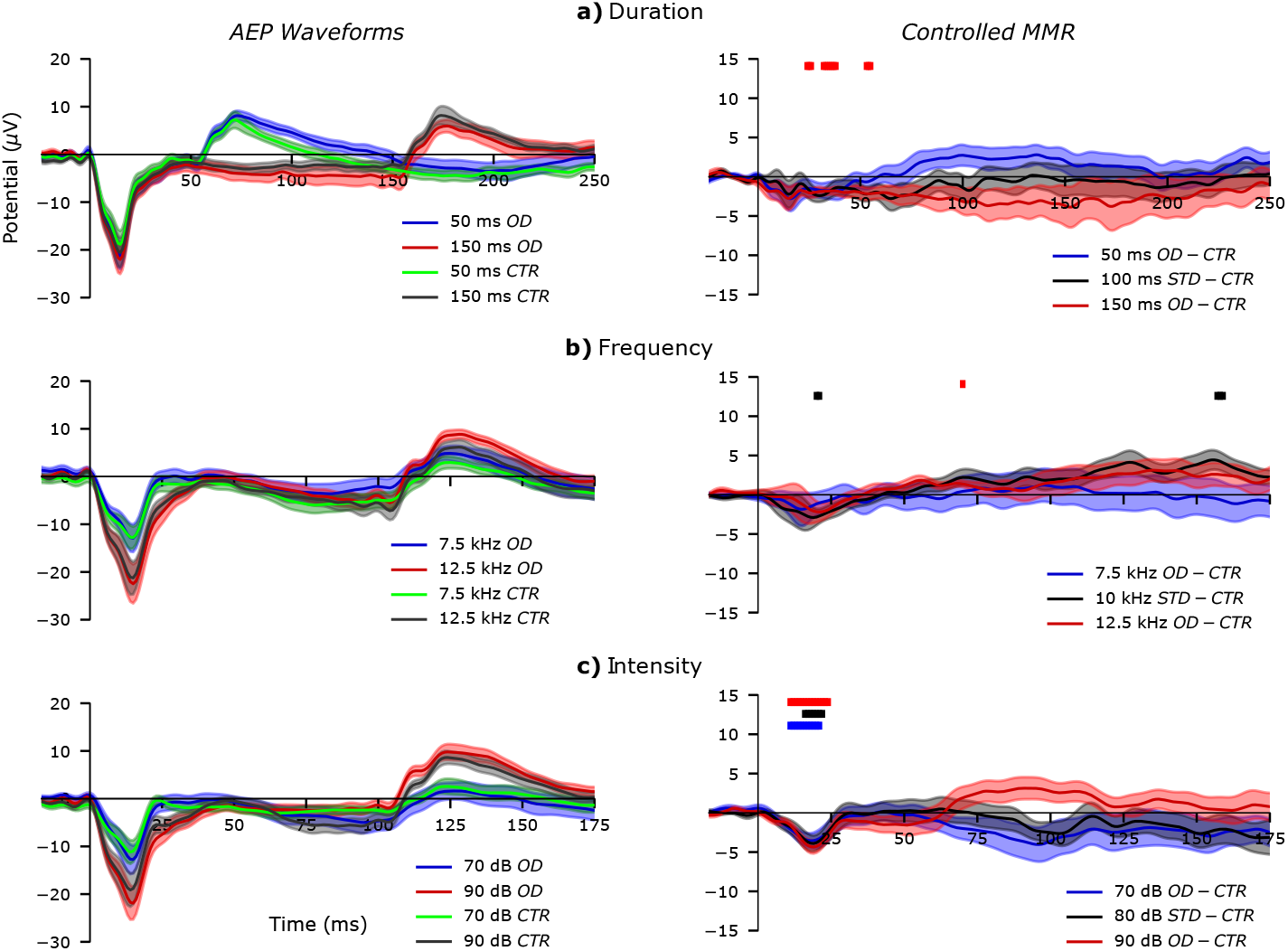
AEP (left) and controlled MMR (right) waveforms from oddball and many-standards paradigms in urethane-anaesthetised mice. Grand-average (n = 14) waveforms ±sem are plotted. a) Duration-varying paradigm waveforms: 50 ms oddball (OD) and control (CTR), 150 ms OD and CTR, 50 ms OD – CTR, 100 ms standard (STD) – CTR, and 150 ms OD – CTR. b) Frequency-varying paradigm waveforms: 7.5 kHz OD and CTR, 12.5 kHz OD and CTR, 7.5 kHz OD – CTR, 10 kHz STD – CTR, and 12.5 kHz OD – CTR. c) Intensity-varying paradigm waveforms: 70 dB OD and CTR, 90 dB OD and CTR, 70 dB OD – CTR, 80 dB STD – CTR, and 90 dB OD – CTR. Note the different timescale for duration-varying waveforms. Statistically significant differences (p < 0.05) are annotated respectively with solid coloured bars.

### 3.2. Conscious Group

The AEP waveforms from duration, frequency, intensity and ISI many-standards paradigms in conscious mice are plotted in Figure 5. Auditory-evoked potential *N*1, *P*1 and *P*_*off*_ peaks are observed, with evident effects from different physical parameters of stimuli. Stimulus duration determined *P*_*off*_ peak latency [F_9,190_ = 8.08, p < 0.0001], quantified from isolated offset response waveforms (O’Reilly, 2019b). Stimulus frequency had a significant relationship with *N*1 peak amplitude [F_9,190_ = 3.00, p = 0.0023] and its effect on *P*1 was nearing the threshold for statistical significance [F_9,190_ = 1.82, p = 0.0668]. Similarly, stimulus intensity had statistically significant relationships with *N*1 [F_9,190_ = 5.53, p < 0.0001] and *P*1 [F_9,190_ = 5.60, p < 0.0001] peak amplitudes. Inter-stimulus interval also had statistically significant relationships with both *N*1 [F_9,190_ = 1.97, p = 0.0450] and *P*1 [F_9,190_ = 4.14, p < 0.0001] peak amplitudes.

**Figure 5:**
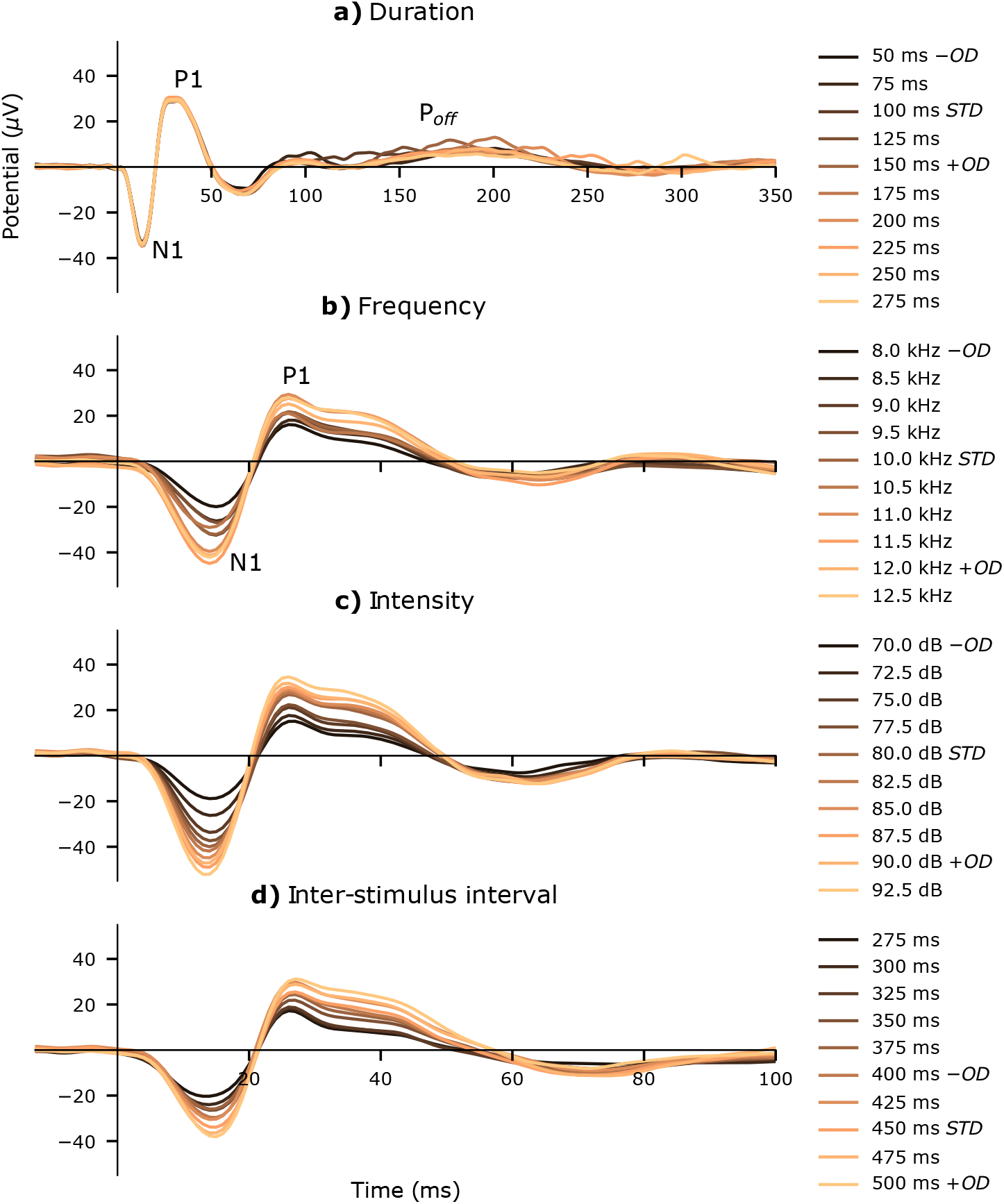
AEP waveforms from many-standards paradigms in conscious mice. This is grand-average data (n = 20). a) Duration-varying stimuli. b) Frequency-varying stimuli. c) Intensity-varying stimuli. d) Inter-stimulus interval (ISI)-varying stimuli. Three components of the AEP are identified; a negative amplitude onset response (*N*1), positive amplitude onset response (*P*1), and positive amplitude offset response (*P*_*off*_). Oddball (+OD and −OD) and standard (STD) stimuli from the oddball paradigms are labelled in the legend. Note the different timescale for duration many-standards control waveforms.

Classical MMR waveforms from duration, frequency, intensity and ISI oddball paradigms in conscious mice are plotted in Figure 6. Prominent negative peak amplitudes in duration MMR waveforms at 125 ms are caused by subtraction of the standard stimulus-off response, and positive peaks at 75 ms and 175 ms are caused by *P*_*off*_ of each respective oddball; although these did not reach the threshold for statistical significance with the tests applied. The longer duration oddball produces greater positive peak amplitude in the MMR, potentially indicating why longer duration oddballs are reported to elicit a greater MMR (Lipponen et al., 2019; Umbricht et al., 2005). Viewing the duration many-standards AEP waveforms (Figure 5a), it may be noted that off-responses from shorter duration stimuli are difficult to distinguish from the larger amplitude deflections. Additionally, offset potentials and difference waveforms in Figure 6a appear to show lower amplitude peaks for the shortest duration stimuli. This is perhaps due to the much larger amplitude *N*1 and P2 deflections occurring during this latency range. However this relationship between offset response peak amplitude and stimulus duration was not found to be statistically significant. Mismatch responses from frequency, intensity and ISI oddballs exhibit similar waveform morphology. Each of these properties of sound has a proportional relationship with *N*1-*P*1 peak amplitudes (Figure 5b-d), and accordingly, increasing and decreasing oddball stimuli tended to generate opposite polarity deflections in their respective difference waveforms. Oddball and many-standards control paradigm waveforms are plotted alongside each other in Figure 7. Controlled MMR waveforms highlight changes that similarly influence oddball and standard waveforms; reflected in an alternating polarity waveform that settles by 100 ms post stimulus onset. These may be interpreted as auditory habituation, or a form of general adaptation, occurring across presentation of subsequent auditory paradigms (oddball followed by many-standards control). The lack of substantial difference between controlled MMR and controlled standard waveforms indicates that neither memory- nor adaptation-based components predominate their morphologies.

**Figure 6:**
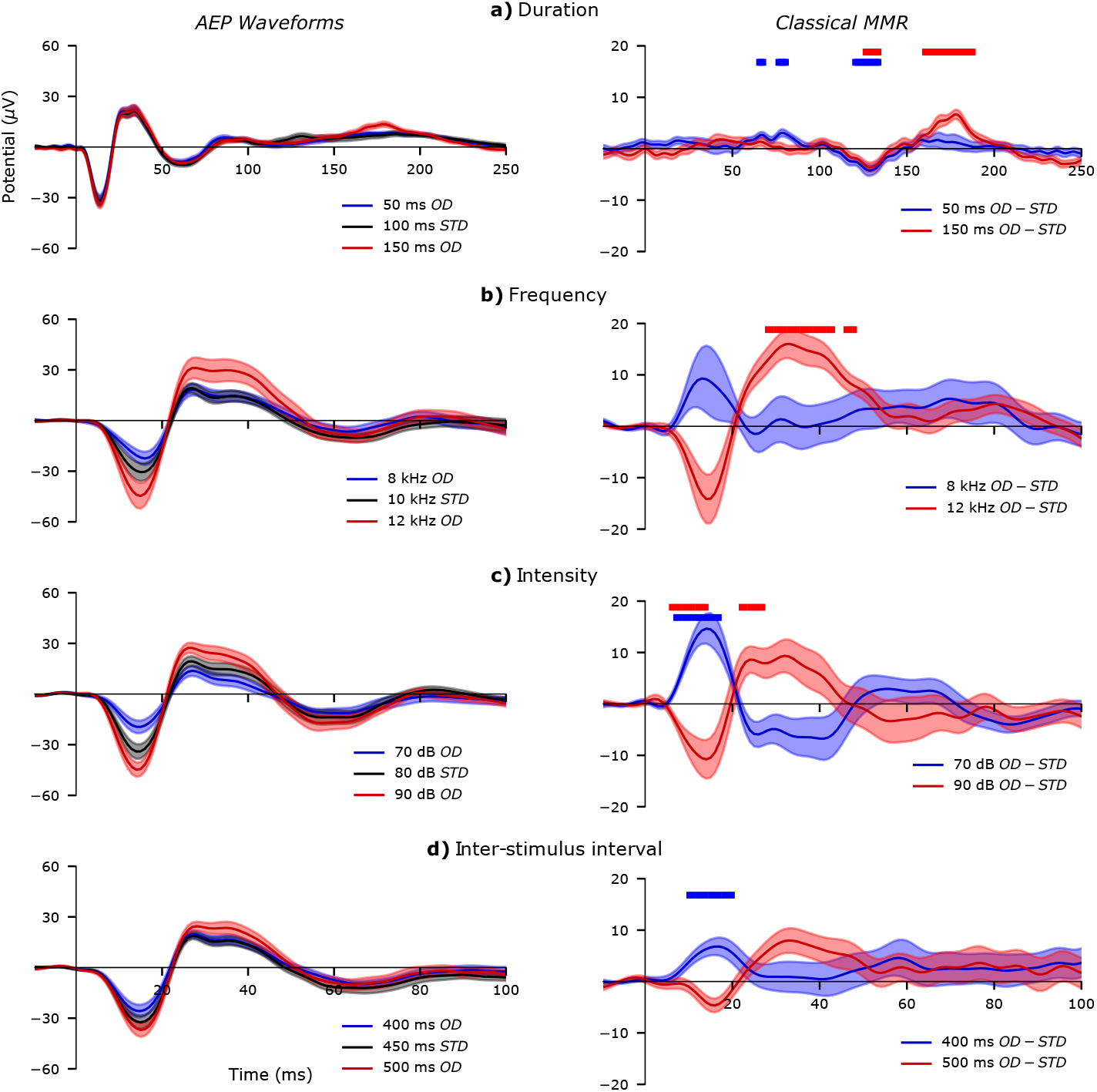
AEP (left) and classical MMR (right) waveforms from oddball paradigms in conscious mice. This is grand-average data (n = 20). a) Duration oddballs minus 100 ms standard (50 ms OD – STD and 150 ms OD – STD). b) Frequency oddballs minus 10 kHz standard (8 kHz OD – STD and 12 kHz OD – STD). c) Intensity oddballs minus 80 dB standard (70 dB OD – STD and 90 dB OD – STD). d) Inter-stimulus interval (ISI) oddballs minus 450 ms standard (400 ms OD – STD and 500 ms OD – STD). Note the different timescale for duration oddball waveforms. Time points that reached the threshold to reject the null hypothesis (p < 0.05) are highlighted with coloured blocks. Shaded regions represent sem.

**Figure 7:**
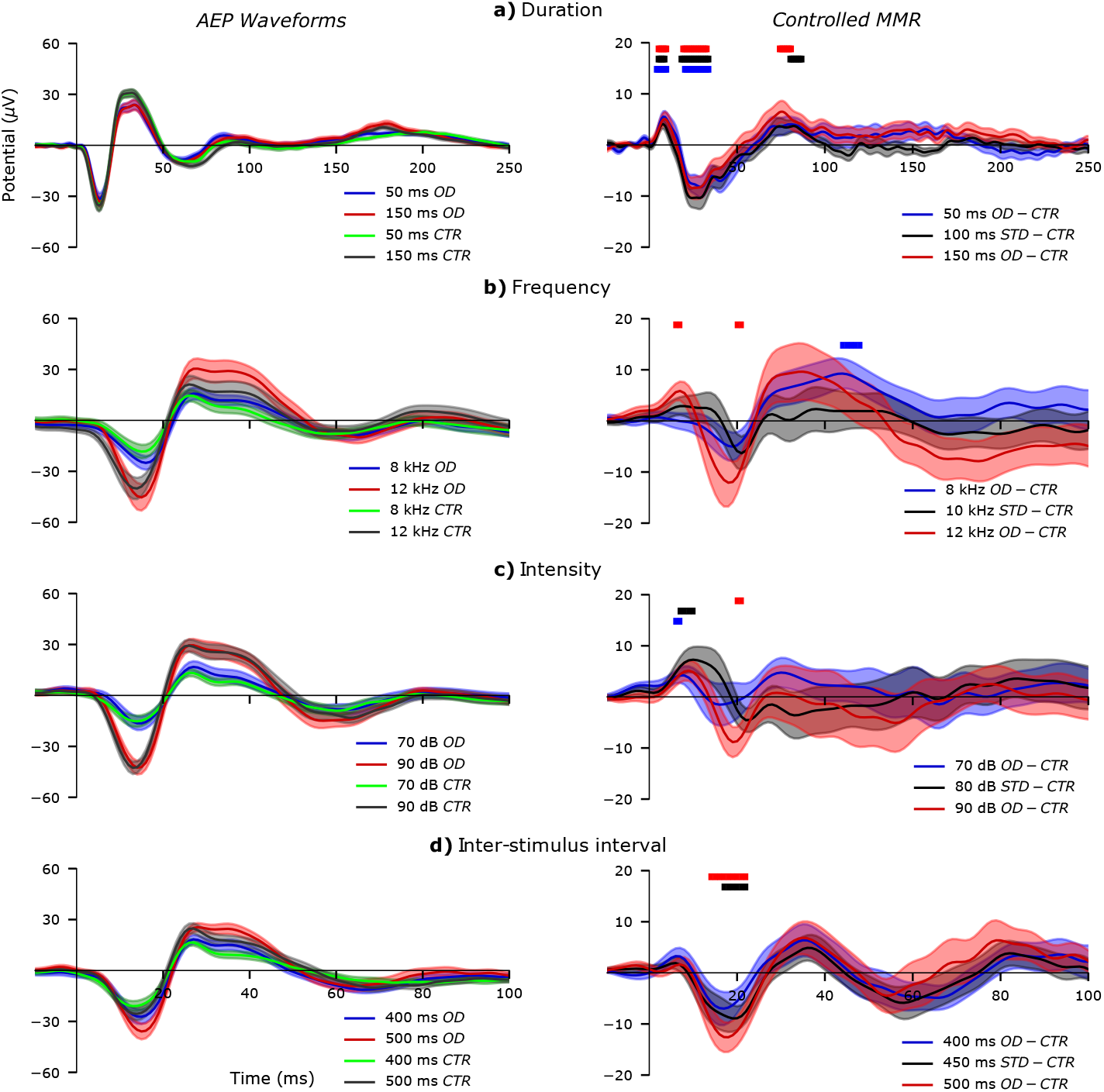
AEP (left) and controlled MMR (right) waveforms from oddball and many-standards paradigms in conscious mice. Grand-average (n = 20) waveforms ±sem are plotted. a) Duration-varying paradigm waveforms: 50 ms oddball (OD) and control (CTR), 150 ms OD and CTR, 50 ms OD – CTR, 100 ms standard (STD) – CTR, and 150 ms OD – CTR. b) Frequency-varying paradigm waveforms: 8 kHz OD and CTR, 12 kHz OD and CTR, 8 kHz OD – CTR, 10 kHz STD – CTR, and 12 kHz OD – CTR. c) Intensity-varying paradigm waveforms: 70 dB OD and CTR, 90 dB OD and CTR, 70 dB OD – CTR, 80 dB STD – CTR, and 90 dB OD – CTR. Note the different timescale for duration-varying waveforms. Statistical tests that exceeded the threshold for statistical significance (p < 0.05) are denoted with coloured blocks.

## 4. Discussion

### 4.1. Anaesthetised and Conscious States

The discussion begins with a qualitative comparison of AEPs from conscious and anaesthetised groups. Stimulus-on (*N*1) and stimulus-off (*P*_*off*_) peaks were common to both urethane-anaesthetised and conscious states, whereas *P*1 and later deflections were only observed from conscious animals. This agrees with previous studies, in which anaesthetised rodents tend to exhibit a less dynamic evoked potential (Harms et al., 2014; Kurkela et al., 2018; Nakamura et al., 2011). One interpretation of this finding is that the neurophysiological mechanisms responsible for generating *P*1 and subsequent deflections are blocked by anaesthetics. These components may signify aspects of consciousness underpinned by neurotransmitter systems disrupted by urethane; for example, GABA, glycine, N-methyl-D-aspartate (NMDA), *α*-amino-3-hydroxy-5-methyl-4-isoxazolepropionic acid (AMPA), or acetylcholine (ACh) receptor-mediated signalling (Hara and Harris, 2002). Based on this, it is plausible to suggest that alternative anaesthetics, with different mechanisms of action to urethane, might produce distinct AEP morphologies. Differences in the AEP between urethane-anaesthetised and conscious mice influence their respective MMR waveforms. This notwithstanding, they also share some similarities. A prominent deflection is present during the *N*1 latency range in response to frequency, intensity and ISI oddballs from animals in both states. Furthermore, classical duration MMR waveforms from both groups are principally formed by offset responses. It may be argued that these correspondences are comparable with human MMN, which is also observed in conscious and unconscious states (Fischer et al., 1999, 2000; Heinke et al., 2004; Kane et al., 1996).

### 4.2. Duration MMR

Stimulus duration is responsible for offset response peak latency (O’Reilly, 2019b). This appears to be instrumental in shaping classical duration MMR difference waveforms; positive peak latency was determined by oddball offset response and negative peak latency determined by the standard offset response (according to the *oddball – standard* computation). This may explain prior findings in conscious mice which found a MMR which varied depending on the duration of standard and oddball stimuli (Umbricht et al., 2005). This also appears to resemble findings from human duration MMN (Colin et al., 2009; Takegata et al., 2008). In a previous study examining duration and frequency deviants in rats, stimulus-off responses were observed from duration-varying stimuli when animals were anaesthetised, but not when they were conscious (Nakamura et al., 2011); for statistical analysis, the investigators average together duration and frequency oddball waveforms, and possible influence of stimulus-off responses is not addressed. Combining duration and frequency oddball-evoked waveforms in this way might be inappropriate, given the apparent distinctions between them (Erickson et al., 2016; Michie et al., 2000; Todd et al., 2008; Umbricht et al., 2005; Umbricht and Krljes, 2005). A more recent study investigating duration deviance in urethane-anaesthetised mice also found a greater amplitude response to longer duration stimuli, although concluded that this may be explained by adaptation (Lipponen et al., 2019). This study used syllable sounds, which are spectrally more complex than monophonic pure-tones, partially obscuring comparison of findings. In the present study, offset potentials were more explicit than previous reports. This is likely due to the rapid stimulus fall times, combined with other factors such as experimental design, electrode placement, animal movement, background acoustics, and electrical interference. For example, a relatively high acoustic signal-to-noise ratio (>25 dB) may have accentuated off-set responses, enhancing their visibility from anaesthetised and conscious animals (Baltzell and Billings, 2014; O’Reilly, 2019b). Stimulus duration, SPL and fall-time also influence cortical activity at sound cessation (Jung et al., 2013; Takahashi et al., 2004). Offset responses are known to occur throughout the auditory system at levels of the brainstem (Henry, 1985), inferior colliculus (Brand et al., 2017), thalamus (He, 2002) and auditory cortex (Qin et al., 2007) in animals/rodents under anaesthesia. These are also observed in EEG recordings from conscious humans (Hari et al., 1987; Hillyard and Picton, 1978), although are only tenuously linked with duration MMN (Jacobsen and Schröger, 2003).

Both onset and offset responses are reactions to abrupt changes in the auditory environment, although their physiological underpinnings remain to be fully elucidated (Yamashiro et al., 2009). It has been proposed that auditory stimulus-off responses reflect post-inhibitory rebound following auditory stimulation (Kuwada and Batra, 1999; Phillips et al., 2002; Takahashi et al., 2004). Considering this interpretation, the offset response may reflect the collective activity of inhibitory neurons acting to ‘quiet’ excitatory neurons in the auditory cortex responding to feedthrough of auditory stimulation from the thalamus. This may be why the onset response peaks before the AEP signal returns towards baseline; when auditory stimulation is removed an overshoot of inhibitory activity occurs, observed here as *P*_*off*_. However, it is argued that both onset and offset responses in the auditory cortex involve separate afferent pathways, suggesting that both are independently driven processes (Scholl et al., 2010). Some research in animals indicates that neurons in the auditory system are fine-tuned to respond to specific duration stimuli by firing an action potential. Short-, long- and band-duration tuned neurons are reportedly found in the auditory cortex, as are neurons which fire at tone offset (Galazyuk and Feng, 1997; He et al., 1997). In the mouse inferior colliculus, duration-tuning properties and stimulus-off triggered neurons have also been reported (Brand et al., 2017). In the primary auditory cortex of cats, offset-specific neurons have been classified and compared with onset neurons, finding that both are actively triggered (Qin et al., 2007), rather than the off response arising from an inhibition-rebound effect. Implicitly, these effects cannot be disentangled, because stimulus offset cannot occur without first having an onset. Given the large number neurons in the auditory system, and current limitations of recording techniques, the relationship between onset and offset responses must remain an open question. Nevertheless, the findings of this study directly implicate the stimulus-off response as a key determinant of classical duration MMR in mice. This concept has been touched upon previously in the human MMN literature (Jacobsen and Schröger, 2003). Furthering our understanding of this neurophysiological process may be beneficial for interpreting AEP deficits in patients with schizophrenia and genetically susceptible individuals (Colin et al., 2009; Shelley et al., 1991; Takegata et al., 2008). Examination of controlled duration MMR waveforms only illustrated auditory habituation, common to both oddball and standard responses, providing no support for any other mechanism underlying the duration MMR in anaesthetised or conscious mice.

### 4.3. Frequency, Intensity, and ISI MMR

Stimulus frequency, intensity and ISI manipulations similarly influenced *N*1, *P*1 and *P*_*off*_ features of the mouse AEP. This is also true for human *N*100 and *P*200 responses (Picton et al., 1977), potentially suggesting that these features may reflect comparable neurophysiological substrates shared between species. These sensitivities caused deflections in each respective classical MMR difference waveform over the latency ranges of these AEP peaks. Opposite polarity deflections were observed for decreasing versus increasing deviances. The magnitude of these MMR waveforms inherently varies with oddball distance from the standard, which is also true for the human MMN (Pakarinen et al., 2007). However, mouse MMR deflections may be wholly ascribed to the physical characteristics of stimuli themselves, and implicit differences in the resulting magnitude of evoked electrophysiological activity. This does not clearly reflect sensory-memory or adaptation mechanisms thought to underlie the human MMN, and rather indicates physical modulation of obligatory AEP components. Inspection of controlled standard and controlled MMR waveforms revealed uniform auditory response habituation, without evidence for memory or adaptation components elicited by the oddball condition. Data from rats also suggests that physical properties of stimuli are instrumental in shaping the resulting frequency MMR (Ruusuvirta et al., 2015). This may question the utility of rodents for modelling human MMN. Conversely, the mechanisms underlying human MMN to different physical properties of sound are not completely understood, and may require further clarification to eliminate this as a possibility. Inclusion of appropriate control paradigms in future human studies is therefore highly recommended.

Ascending frequency oddball stimuli in conscious mice produced larger amplitude classical MMR over the P2 latency range. This may reflect non-linearity of the mouse auditory system, which is reported to exhibit tone preference towards ≈14 to 18 kHz (Heffner and Heffner, 2007). The findings here may indicate that AEP amplitudes increase with hearing sensitivity. To examine whether this is true, stimulus frequencies exceeding the range of greatest hearing sensitivity may be applied to test whether a symmetrical decay in amplitude occurs with increasing tone frequencies; although this would require specialist audio equipment, which was not accessible for the present study. An argument may be made that the oddball frequencies were too far removed from the standard (by ±2.5 kHz and ±2 kHz in anaesthetised and conscious groups, respectively), thus influenced by variable hearing thresholds, impeding comparison with human studies. This degree of deviance between standard and oddball stimuli accounts for approximately ±2 to 2.5% of the mouse hearing range; comparable with relative frequency deviances previously employed in oddball paradigms for human subjects (Michie et al., 2000). The frequencies and deviances used here are also within the usual range, if slightly towards the higher end, of those typically applied in rodent MMR studies (Harms et al., 2014; Kurkela et al., 2018; Nakamura et al., 2011; Polterovich et al., 2018; Shiramatsu et al., 2013). Furthermore, the degree of deviance between standard and oddball stimuli has a well-documented effect, which is reportedly to enlarge the MMN amplitude (Colin et al., 2009; May et al., 1999; Shelley et al., 1991; Takegata et al., 2008). This appears to emphasise the fact, because different physical properties of sound have a proportional relationship with AEP amplitudes, that increased separation between the standard and oddball stimuli frequencies inherently produce greater amplitude difference waveforms.

Effects of stimulus intensity/SPL on the AEP have been studied extensively in both humans and animals. Loudness dependence of the AEP (LDAEP), comparable with observations from this study, has been proposed to reflect serotonergic, dopaminergic and glutamatergic neurotransmission in the primary auditory cortex (Hegerl and Juckel, 1993; Nathan et al., 2005; O’Neill et al., 2008). This may demonstrate characteristic firing patterns of intensity-tuned, non-monotonic and monotonic neurons (Phillips and Irvine, 1981). Altered LDAEP is associated with a range of neuropsychiatric diseases, including schizophrenia (Juckel, 2015; Park et al., 2010), which could potentially explain the presence of intensity MMN deficits in patients. Thus, intensity MMR deficits may be due to altered intensity modulation of obligatory AEP components. Inter-stimulus interval modulation of AEP amplitudes has been linked with NMDA receptor signalling, which interestingly is also widely implicated in MMN (Javitt, 2000). To the author’s knowledge no neurotransmitter systems have been linked with frequency modulation of the AEP. There is a relative paucity of published studies investigating intensity or ISI MMRs in rodents for comparison, but the data presented here strongly suggests that these are due to physical sensitivities of the auditory system, not memory or adaptation mechanisms. The use of physically identical control waveforms in rodent studies (Harms et al., 2014; Nakamura et al., 2011) does not appear to be widely shared in human MMN studies (Erickson et al., 2016; Näätänen et al., 2012). The majority of research in humans and animal models follows the conventional approach, subtracting standard from oddball AEPs to generate the MMN/MMR waveform (e.g. Figure 2 and Figure 5), which is somewhat disconcerting given that modulation of AEP amplitudes with different physical parameters is a well-established finding in auditory neuroscience.

### 4.4. Auditory Habituation

Sequential ordering of paradigms illustrated the effect of auditory habituation (Picton et al., 1976) on AEP waveforms (Figure 4 and Figure 7). Amplitudes gradually diminished across paradigms. If this were due to SSA, we would expect to see a differential influence on controlled MMR versus controlled standard responses, because the former represents two conditions with the same presentation rate, while the latter reflects the difference between higher and lower stimulus presentation rates (Nelken, 2014). However, amplitude reductions observed in controlled MMR and controlled standard waveforms were comparable, discounting any overt influence of SSA. Furthermore, lack of substantial differences between controlled MMR and controlled standard waveforms indicates an absence of memory-based component elicited by the oddball. We argue that this approach is more robust than a counterbalanced study design (such as the flip-flop sequence combined with a many-standards control (Harms et al., 2014)), which would distribute auditory habituation between subjects across different paradigms. Differential subject habituation rates or minor imbalances in the ordering of paradigms between subjects could lead to considerably uneven distributions of auditory habituation affecting the data. Perhaps this can explain some of the positive findings from previous rodent MMR studies.

Counterbalancing may be taken for granted as a simple procedural control, but when trying to arrange an odd number of paradigms evenly across all subjects imbalances can easily occur. This may be particularly evident where two sides of the flip-flop oddball paradigm are counterbalanced then followed by another control sequence. This would decrease the probability of finding a significant difference between AEPs from identical stimuli presented in oddball and standard conditions (dismissing the adaptation hypothesis), while increasing the likelihood of observing differences between oddball and control conditions (falsely confirming the memory hypothesis). Few studies we are aware of provide adequate counterbalancing details to fully dissociate this possibility (Harms et al., 2014; Kurkela et al., 2018; Polterovich et al., 2018). The series of waveform comparisons presented here (Figure 1) on sequentially ordered paradigm data facilitate an inspection of physical sensitivity, auditory habituation, sensory-memory, and adaptation mechanisms that may influence difference waveform morphology, while addressing the potential confound of uncontrolled auditory habituation between paradigms. The only phenomena that were clearly observed from these analyses were physical modulation of AEP waveforms and habituation of response amplitudes with repeated stimulation. Thus no evidence was found to objectively support either of the two prevailing hypotheses for MMR generation.

## 5. Conclusion

These findings demonstrate that mismatch response morphology is primarily shaped by physical differences between oddball and standard stimuli. When MMR difference waveforms are computed by the conventional method, these inherent effects can explain all of the observed non-zero amplitudes. This suggests that classical MMR in mice does not reflect a violation of auditory sensory-memory, or adaptation, which are both thought to contribute towards human MMN. Moreover, analysis of controlled MMR and controlled standard waveforms provides no support for either of the existing hypotheses. Overall, this suggests that difference waveforms in mice are fundamentally influenced by physical sensitivity and habituation of the auditory response. Further work should be cognizant of this interpretation and apply appropriate experimental constraints to confirm the nature of difference waveforms derived from the oddball paradigm.

## 6. Materials & Methods

### 6.1. Animals

Laboratory mice (>99.999% C57BL/6J) were used in this study. The anaesthetised group (n = 14) consisted of 9 males and 5 females aged 14 to 17 weeks (mean 15.4). Conscious group (n = 20) consisted of 9 males and 11 females aged 29 to 37 weeks (mean 32.4). These two separate cohorts, which may be considered young and middle-aged adults, respectively, have sufficient hearing capacity for inclusion in this experiment (Ison et al., 2007). Qualitative between-groups analyses are appropriate, given this age disparity. Animal husbandry followed standard guidelines for laboratory mice. All procedures were approved by the Animal Welfare and Ethical Review Board at University of Strathclyde, and performed in accordance with the UK Animals (Scientific Procedures) Act 1986.

### 6.2. Surgery

Urethane-anaesthesia was applied as described elsewhere (Sakata and Harris, 2009), and depth of anaesthesia was confirmed throughout experiments by an absence of normal (tail and toe pinch, eye-blink) reflexes. The conscious experiment group were anaesthetised with isoflurane and oxygen for the duration of aseptic surgery. Skull-screw electrodes (1 mm diameter; Royem Scientific Ltd., Luton, UK) were used to record epidural field potentials. Recording electrodes were implanted bilaterally above the primary auditory cortices (2.2 mm caudal, 3.8 mm lateral, relative to Bregma; Paxinos and Franklin, 2004), with a ground electrode implanted above the cerebellum; as previously described (O’Reilly, 2019b). The conscious group were fitted with a connector (SDL-12-G-10; Samtec, IN, USA), fixed to the skull with dental acrylic, wired to electrodes, providing a conductive path for the amplifiers; after surgery mice recovered for five days before beginning electrophysiological recordings.

### 6.3. Electrophysiology

Mice were placed inside a customised enclosure for electrophysiology recording sessions. This consisted of a Faraday cage within an acoustically-controlled cubicle; the inner walls were coated with 50 mm, corrugated, sound-absorbing foam. Background noise levels were maintained below 55 dB sound pressure level (SPL). Anaesthetised animals were positioned facing a loudspeaker, calibrated to the approximate position of their head using a sound meter (A-weighted). Conscious mice were held in a perforated isolation tube (100/44 mm; length/diameter) positioned between front and rear loudspeakers; the average of two calibrations with the sound meter facing either direction was taken in this case. Mice tended to orient themselves longitudinally within the tube, facing either of the speakers. This provided some control over relative sound source location, which is also thought to induce a mismatch negativity response (Perrin et al., 2018), while also reducing movement-related artifacts. The isolation tube had a narrow cut-out along the top, allowing the amplifier tether to enter. Conscious mice were gradually habituated to the test equipment with increasing familiarisation sessions of 5, 15, and 30 min durations on separate training days before beginning recordings. Epidural field potentials were acquired at a sampling rate of 1 kHz and band-pass filtered from 0.1 to 500 Hz using a tethered amplifier board (Intan Technologies, CA, USA) and stored for post-hoc analysis (open-ephys.org). Stimuli and synchronisation signals were generated in Matlab (Mathworks Inc., MA, USA) and output via a USB i/o device (USB-6255; National Instruments, TX, USA).

### 6.4. Stimulation

Three balanced oddball paradigms, containing both increasing and decreasing deviants, were presented to the anaesthetised group; a separate one for stimuli varying in duration, frequency, and intensity. In addition, the conscious group were presented with an inter-stimulus-interval (ISI) varying oddball paradigm. Inter-stimulus interval is defined as the stimuli offset to onset time. Each oddball paradigm consisted of 800 standards, 100 increasing-oddballs, and 100 decreasing-oddballs; with an ISI of 450 ms (although this was altered in the ISI-varying paradigm). The separation of physical feature variance into different paradigms enabled an examination of each property of sound individually, mitigating against potentially confounding effects of multiple-deviances. An initial sequence of 20 standards was followed by pseudo-random presentation of either increasing or decreasing oddballs, with at least three intervening standards. Standard stimuli were 100 ms, 10 kHz sinusoidal, 80 dB monophonic tones. Duration oddball stimuli varied by ±50 ms; frequency oddball stimuli varied by ±2.5 kHz (anaesthetised group) or ±2 kHz (conscious group); intensity oddball stimuli varied by ±10 dB; and ISI oddballs varied by ±50 ms. All stimuli had instantaneous rise and fall times; this caused no distortion of electrical recordings.

Three many-standards control paradigms were presented to the anaesthetised group; the conscious received four, with the addition of an ISI-varying control. These consisted of 10 different level stimuli, including the standard and both oddballs from the oddball paradigm, presented pseudo-randomly 100 times each, with an ISI of 450 ms (although this was altered in the ISI-varying paradigm). For each version of the many-standards paradigm, the other two physical parameters of stimuli were constant (e.g. in duration-varying paradigms, stimuli frequency and intensity were constant), maintained identical to the standard in the oddball paradigm. The duration many-standards paradigm employed stimuli varying from 50 to 275 ms in 25 ms increments. In the anaesthetised group frequency many-standards stimuli varied from 1.25 to 12.5 kHz in 1.25 kHz increments; in the conscious group these varied from 8 to 12.5 kHz in 500 Hz increments. In the anaesthetised group, intensity many-standards stimuli varied from 60 to 105 dB in 5 dB increments; in the conscious group these varied from 70 to 92.5 dB in 2.5 dB intervals. The adjustments in stimuli frequency and intensity ranges between groups followed observation of very low-amplitude AEPs in response to stimuli at the lower end of these scales in the anaesthetised group. The ISI many-standards control paradigm incorporated 10 different levels of ISI that varied from 275 to 500 ms in 25 ms increments.

The anaesthetised group were presented with auditory paradigms immediately following surgery, in a structured protocol lasting approximately 90 min. Duration, frequency and intensity paradigms were presented sequentially, allowing an examination of the influence of deepening anaesthesia and repeated bouts of auditory stimulation on the elicited auditory responses. Level of anaesthesia was intermittently monitored by ensuring absence of normal palpebral and corneal reflexes; no reacquisitions of consciousness were observed. The conscious group were presented with counterbalanced duration, frequency, intensity, and ISI-varying paradigms on separate test days, with one intervening non-test day between each; sessions lasted approximately 30 min. In both groups, each feature-specific oddball paradigm was followed by its respective control; e.g. the duration oddball paradigm was followed by the duration many-standards control, facilitating inspection of auditory habituation across paradigms.

### 6.5. Data Analysis

Data were processed with a zero phase-shift low-pass filter with cut-off frequency of 100 Hz. Segments were extracted from −100 to +400 ms about a stimulus-onset timestamp. Pre-stimulus average baseline correction was applied and trials containing voltages exceeding ±500 *μ*V were removed. Preliminary analysis revealed no significant hemispheric differences between AEPs, comparable with previous findings (Harms et al., 2014). The channel-averaged AEP was therefore computed for each animal, combining left and right electrode data. Only standards immediately preceding oddballs were included in this computation to maintain a relative balance between the number of standard and oddball sweeps contributing to averages.

Significant non-zero amplitudes from difference waveforms were quantified by statistically evaluating every time point. Any adaptation effects were analysed in the controlled standard MMR (standard from the oddball paradigm minus the same stimulus in the many-standards control). This difference wave reflects the distinction between a repetitive stimulus, presented with 80% probability, thereby dominating short-term sensory-memory, and a physically identical stimulus presented at 10% probability, without a predictable recent history of auditory stimulation. Neither of these conditions is regarded to be in violation of a sensory-memory trace; thus, any significant differences between them may be ascribed to SSA. Moreover, given the order of presentation, waveforms elicited by oddball and control sequences may reflect changes in anaesthetic state and/or habituation of the auditory response over a period of repeated stimulation (Picton et al., 1976). In order to examine whether any specifically memory-based features were elicited by the oddball paradigm, controlled MMR waveforms (oddball minus control) were compared with controlled standard MMR waveforms; wherefore memory-violation may be dissociated from both physical sensitivities, because each evoked response is effectively normalized against that of a physically identical stimulus, and SSA or habituation, reflected in the controlled standard MMR. Wilcoxon signed-rank tests were performed at every time-point from 350 ms post-stimulus onset with alpha set to 0.05; no corrections for multiple comparisons were applied, given that this may impose excessive requirements for achieving statistical significance from this highly sampled data. Regions exceeding the threshold for statistical significance are indicated in the relevant figures.

Auditory evoked potential features quantified from many-standards control wave-forms include stimulus-on components, *N*1 and *P*1 (conscious animals only), and the stimulus-off response, *P*_*off*_. These were detected over 0 to 30 ms post-onset, 20 to 50 ms post-onset, and 0 to 50 ms post-offset measurement windows, respectively. In the conscious group, stimulus-off potentials from duration many-standards paradigm stimuli were isolated by subtracting the mean AEP from each individual AEP, as described previously (O’Reilly, 2019b). The effects of stimulus duration, frequency, intensity, or ISI on these AEP features were tested with a one-way, within-subjects, repeated measures analysis of variance, using many-standards paradigm data. The Bonferroni adjustment was applied to correct the alpha value to 0.005; taking into consideration ten discrete stimulus levels in many-standards paradigms. Shaded waveform error-bars represent standard error of the mean (sem). These analyses were conducted using MNE-Python (Gramfort et al., 2013) and R-studio. The data described in this article are stored in an open-access repository (O’Reilly, 2018).

## 7. Acknowledgements

The doctoral supervision of Professor Judith A. Pratt is much appreciated. This work was supported by funding from the United Kingdom Engineering and Physical Sciences Research Council (EP/F50036X/1).

